# Differential integrin adhesome expression defines human natural killer cell residency and developmental stage

**DOI:** 10.1101/2020.12.01.404806

**Authors:** Everardo Hegewisch Solloa, Seungmae Seo, Bethany L. Mundy-Bosse, Anjali Mishra, Erik Waldman, Sarah Maurrasse, Eli Grunstein, Thomas J. Connors, Aharon G. Freud, Emily M. Mace

## Abstract

Natural killer (NK) cells are innate immune cells that reside within tissue and circulate in peripheral blood. As such, they interact with a variety of complex microenvironments, yet how NK cells engage with these varied microenvironments is not well documented. The integrin adhesome represents a molecular network of defined and predicted integrin-mediated signaling interactions. Here, we define the integrin adhesome expression profile of NK cells from tonsil, peripheral blood and those derived from hematopoietic precursors through stromal cell coculture systems. We report that the site of cell isolation and NK cell developmental stage dictate differences in expression of adhesome associated genes and proteins. Furthermore, we define differences in cortical actin content associated with differential expression of actin regulating proteins, suggesting that differences in adhesome expression are associated with differences in cortical actin homeostasis. Together, these data provide new understanding into the diversity of human NK cell populations and how they engage with their microenvironment.

## Introduction

Human natural killer (NK) cells are commonly defined as CD56^+^CD3^−^ cytotoxic innate lymphocytes and they play a critical role in identifying and killing virally infected or malignant cells. The importance of NK cells in the control of viral infections is underscored by the clinical course of patients with NK cell deficiencies who experience severe and often life-threatening viral infections (1–3). In addition to circulating NK cells found in peripheral blood, tissue resident NK cell populations are present in organs including liver, lung, spleen, bone marrow, and secondary lymphoid tissue, where they serve unique cytotoxic and regulatory functions (4, 5). NK cell maturation is marked by the progressive gain of NK cell-associated receptors and functions, and NK cell developmental subsets can be defined as stages 1-6, which are stages of maturation that represent unique cell phenotypes and lineage potentials (6–12). The stage 4 NK cell subset can be further delineated to stages 4A and 4B by expression of NKp80 at stage 4B (13). The predominant NK cell subsets in peripheral blood are stages 4B, 5 and 6, often defined as CD56^bright^, CD56^dim^ and terminally mature CD56^dim^ respectively, however circulating NK cell and innate lymphoid cell precursors are also found at low frequencies (7–9, 13, 14). CD34^+^ NK progenitor cells are thought to enter circulation from bone marrow and subsequently seed peripheral sites to continue NK maturation; functionally mature NK cells then re-enter circulation in peripheral blood as stage 4B or stage 5 effectors (7). As such, stage 5 cells predominate in circulation, express perforin and granzymes at baseline, and are considered poised for cytolytic function following recruitment to sites of infection or inflammation. Tissue resident NK cells are generally considered to be stage 4 (CD56^bright^) NK cells, yet have distinct phenotypes from peripheral blood stage 4 cells and are thought to perform more regulatory functions (4, 10). The differential expression of chemokine receptors and adhesion molecules on NK cells in peripheral blood and tissue have been implicated in the mechanisms that mediate tissue localization and homing of distinct NK cell subsets (4, 5).

Integrins act as bidirectional signaling hubs between cellular machinery, namely the actin cytoskeleton, and the extracellular microenvironment. As such integrins play particularly critical roles in lymphocyte activation, immune synapse formation and B, T, and NK cell development (15–20). Integrin function is finely tuned and can be regulated by changes in expression, localization and affinity, and their specificity for ligand is dictated in part by the pairing of 18 alpha subunits and 8 beta subunits to form at least 24 unique non-covalently linked obligate heterodimers. NK cells express leukocyte-specific β2 integrins that mediate cell-cell interactions, as well as those that are more broadly expressed and facilitate cell-matrix interactions, including β1 and β7 integrins (21–27). Previous studies have shown that VLA-4 (α4β1, CD49d/CD29) and VLA-5 (α5β1, CD49e/CD29) heterodimers are expressed by NK cells found in peripheral blood and mediate NK cell adhesion to fibronectin (27). LFA-1, the αLβ2 heterodimer (CD11a/CD18), and Mac-1, the αMβ2 heterodimer (CD11b/CD18), mediate NK cell adhesion to ICAM-1 on target cells, leading to signaling and actin remodeling which stabilize the immunological synapse of NK cells (24–26). Furthermore, a subpopulation of NK cells in human peripheral blood are detected with LFA-1 in a partially activated conformation, suggesting that conformational regulation, in addition to expression, is a feature of integrin phenotypes (25). Differences in tissue residency and function between NK cell developmental subsets suggest that distinct integrin repertoires mediate critical functions in lymphocyte development, activation and migration. However, the expression patterns of integrins and their associated signaling networks from NK cell subsets have not been comprehensively described.

The integrin adhesome is the in silico molecular catalogue of signaling protein interactions that occur in response to integrin engagement with ligands, including extracellular matrix (ECM) components and cell adhesion molecules (28–32). While integrin adhesion complexes are more dynamic in lymphocytes than in larger, slower-moving cells, components of the integrin adhesome are conserved and mediate critical functions in lymphocyte development, trafficking and activation (28–32). Using RNA sequencing (RNA-Seq) and phenotypic analyses, we sought to define adhesome expression within human NK developmental subsets from tonsil, a known site of NK cell development, peripheral blood, and in vitro-derived NK cells. We found that differences in the site of isolation and NK cell developmental intermediates are associated with unique profiles of integrins, actin regulators and other signaling intermediates. We additionally revealed significant differences in the density of cortical actin between NK cell developmental subsets in peripheral blood, thus linking integrin-mediated actin signaling networks with inherent differences in actin networks.

## Results

### Tonsil and peripheral blood NK developmental subsets have unique profiles of adhesome gene expression

While differences in integrin expression in NK cell developmental subsets have been previously reported (5, 25, 27), integrin adhesome expression in human NK cells has not been extensively catalogued. Using flow cytometry to sort phenotypically equivalent NK cell developmental subsets, we isolated stage 4B, 5, and 6 NK cells from peripheral blood (PB) and stage 3, 4A, 4B, and 5 NK cells from tonsils obtained from routine tonsillectomies performed on healthy children (9, 10, 13, 33). As CD57^+^CD56^dim^ stage 6 NK cells are found at very low frequencies in tonsils from healthy individuals, they were not included in our experimental design (Supp. Fig. 1). We pooled RNA from 12 donors into 3 technical replicates and performed bulk RNA-Seq. Principal component analysis (PCA) using the 18,475 genes detected by RNA-Seq revealed that both tissue residency and developmental stage are determinants of unique gene expression profiles (Fig. 1A). This analysis revealed that the overall gene expression is more significantly affected by tissue specificity (PC1: 57% of variance) than the developmental stages (PC2: 12% of variance). As such, when phenotypically equivalent stage 4B and 5 cells from peripheral blood or tissue were compared, the two subsets from the same tissue clustered more closely than the two subsets of the same developmental stage (Fig. 1A, Supp. Fig. 2). To further understand the genes contributing to the separation observed between PB and tonsil NK cells, we plotted the 18,475 genes used for PCA by their PC1 loading weights (Fig. 1A). Genes associated with the generally immature phenotype of tonsil NK cells, such as *GZMK, KLRC1*, and *IL7R*, were primarily represented by negative weights in PC1, consistent with the clustering of tonsil subsets with negative PC1 values (Fig. 1B). In contrast, peripheral blood NK cells were marked by positive PC1 weights of genes related to NK cytotoxicity, activation, and KIR receptors (*B3GAT1, KLRB1, KIR3DL2, KIR2DL1*, and *IFNGR1*) (Fig. 1B). This distribution of gene sets across PC1 suggested that the differences between the transcriptome of tonsil and peripheral blood NK cell subsets also reflected differences in the predominant stages of NK cell maturation found at these sites. In addition, we found cytoskeleton-associated proteins were highly contributing to PC1, suggesting that these are among the genes significantly driving the differences between PB and tonsil NK cell subsets. Specifically, we noted integrins including *ITGAD, ITGAE*, and *ITGA1* within the genes with the lowest PC1 weights (<0.01), while other integrins and actin regulator proteins, such as *ITGB2, PXN* and *ARF1*, were within the genes with the highest PC1 weights (>0.01, Fig. 1B). To better understand how integrins were differentially expressed between tissue sites and developmental subsets, we identified the distribution of 229 consensus adhesome genes that have been described previously (30–32) (Supp. Table 1) across PC1 weights (Fig. 1B). Adhesome genes were significantly enriched in the negative end of PC1 (22 adhesome genes in the lowest 5%; Fig. 1B). Further, GO pathway analysis identified pathways associated with cell migration, including regulation of T cell migration (p-value=8.86E-04), dendritic cell migration and positive regulation of cell migration, that were associated with genes with low PC loading weights (Supp. Table 2). These observations suggest that genes which are related to integrin-mediated adhesion and cytoskeletal remodeling are in part driving transcriptional differences between tonsil and PB NK cell subsets. The integrin adhesome network has been described primarily in non-lymphocyte cells (30–32) yet our observations of PC1 weights indicated that distinct patterns of adhesome gene expression are also important in NK cell subsets. To further understand how adhesome genes specifically drive NK cell heterogeneity, we performed PCA only with the 229 consensus adhesome genes (30). Similar to the whole transcriptome PCA (Fig. 1A), PC1 separated tonsil NK cells from PB NK cells, whereas PC2 revealed a tonsil specific progression of adhesome gene expression, suggesting that tonsil NK cells undergo additional changes through development (Fig. 1C). Together, these data and differential gene expression analysis demonstrate that genes associated with the integrin adhesome are differentially regulated between both developmentally and spatially distinct NK cell subsets (Supp. Fig. 3). Further, these findings suggest that adhesome gene expression changes are integral parts of NK cell maturation and their adaptation to new environments.

**Fig. 1.**
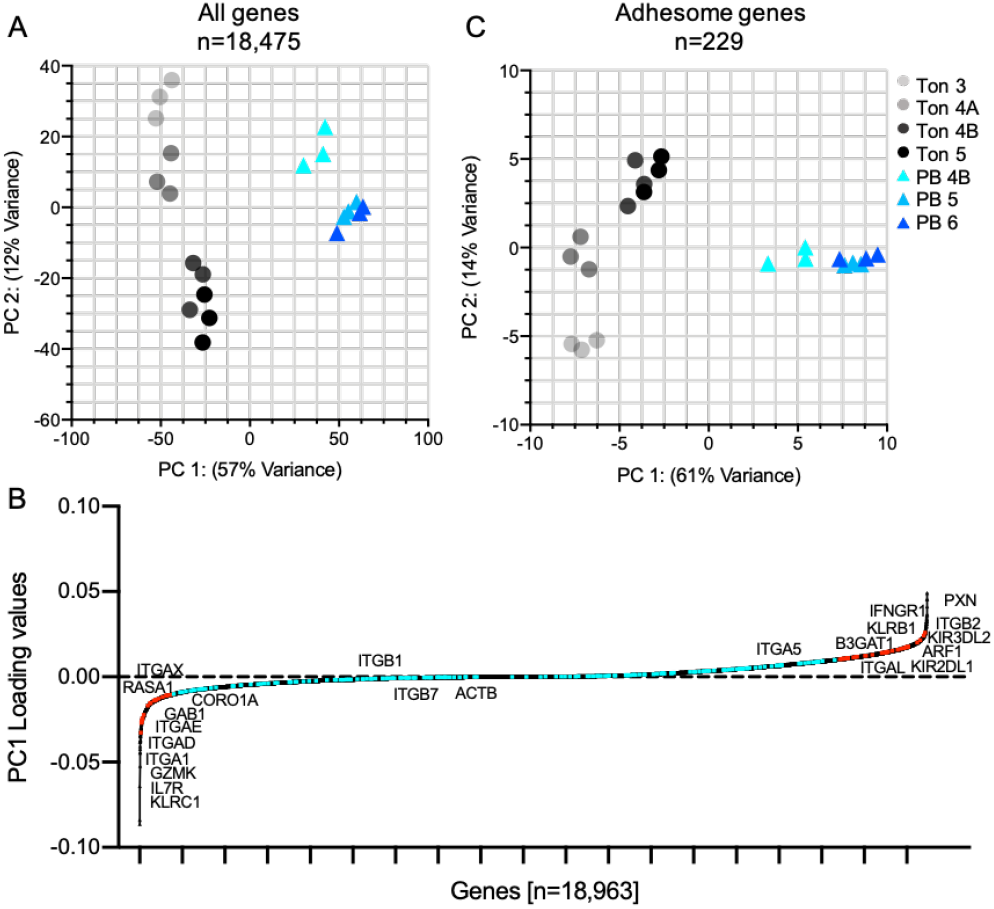
Tonsil and peripheral blood NK cell subsets have unique profiles of adhesome gene expression that reflect distinct developmental trajectories. Bulk RNA sequencing was performed on sorted primary human NK cell developmental intermediates from healthy peripheral blood and tonsil donors as described in Methods. A) PCA of gene expression of NK developmental stages 3, 4A, 4B and 5 isolated from tonsil, and stages 4B, 5 and 6 from peripheral blood. Ton, tonsil; PB, peripheral blood; number represents NK developmental stage. Each data point represents a single technical replicate (n=3) pooled from 12 biological donors. B) 18,475 genes ordered and plotted by PC1 loading weights. 229 adhesome genes are highlighted; red points represent 56 adhesome genes with PC1 weights greater than 0.01 or less than −0.01; turquoise points are adhesome genes with PC1 weight values between −0.01 and 0.01. Labeled points represent relevant adhesome and NK cell associated genes. C) PCA of peripheral blood and tonsil NK subset expression of 229 genes following filtering of whole transcriptome on integrin adhesome genes (Supplemental Table 1).

### Distinct patterns of adhesome gene expression

To further define developmental and tissue residency signatures of tonsil and PB NK cell adhesome genes, we identified 5 clusters (clusters A-E) of adhesome genes with K mean clustering (Fig. 2A, B). Cluster A contain genes that are highly expressed in PB NK cells compared to tonsil NK cells. Cluster A includes β1 integrin partners *ITGA4* (CD49d), *ITGA5* (CD49e), the actin regulator *RAC1*, and *PALLD*, a component of actin microfilaments which functions as an actin stabilizer (Fig. 2B, C). Similar to cluster A, cluster B includes genes that are highly expressed in PB NK cells, but an important distinction is that cluster B genes are upregulated with maturation. Leukocyte-specific integrins including *ITGAL, ITGAM, ITGB2*, and calpain 2 (*CAPN2*), which functions in focal adhesion disassembly (34), are in this cluster. In addition, cluster B reveals that mature PB stage 5 and 6 NK cells upregulate actin and cytoskeletal regulatory pro-teins like *ABI3, ARF1, ARPC2, PXN* and *PI3KCA* (Fig. 2B, C). Cluster C is comprised of genes that are transiently expressed between tonsil stages 4B and 5, and PB stage 5. This includes genes related to the regulation of the cytoskeleton such as *PTK2, GAB1, CORO1A*, and *ITGAX*. Together, genes in clusters A-C show that PB NK cells are characterized by upregulation of actin regulatory proteins and leukocyte associated integrins relative to their tonsil counterparts. Tonsil NK cell subsets were predominantly defined by their preferential expression of adhesome genes belonging to clusters D and E (Fig 2A, B). Cluster D distinguishes tonsil stage 3 and 4A NK cells from the more mature tonsil stage 4B and 5 as well as PB stages 4B-6 (Fig. 2A, B). The genes that constitute cluster D include *ITGB7*, signaling proteins *PRKCA, PAK1*, and *ADAM12*, a disintegrin and metallopeptidase involved in cell migration, proliferation and invasion, and *LRP1*, which is predicted to regulate cell migration through Rho GTPases (35, 36) (Fig. 2B, C). Cluster E defines tonsil NK cells from PB NK cells (Fig. 2A) and is made up of adhesome genes associated with tissue residency. These genes include integrins *ITGA1* and *ITGAE* which encode for CD49a and CD103 respectively and are specifically associated with NK cell tissue residency (21–23) (Fig. 2B, C). Cluster E also reveals that tonsil NK cells preferentially express migration-associated signaling proteins *RASA1, SRC, TIAM1, HSPB1* and *VIM* relative to PB NK cells (Fig. 2B, C). Finally, layilin (*LAYN*), which binds to talin and localizes to membrane ruffles, and neuropilin (*NRP1*), a transmembrane glycoprotein that has not been previously described to play a role in NK cell migration (37, 38) are also found in cluster E and are upregulated by tonsil NK cells relative to PB NK cells (Fig. 2B). Taken together, K means clustering of adhesome genes reveals the unique gene expression signatures in tonsil and PB NK cell subsets that are associated with both tissue residency and developmental stage.

**Fig. 2.**
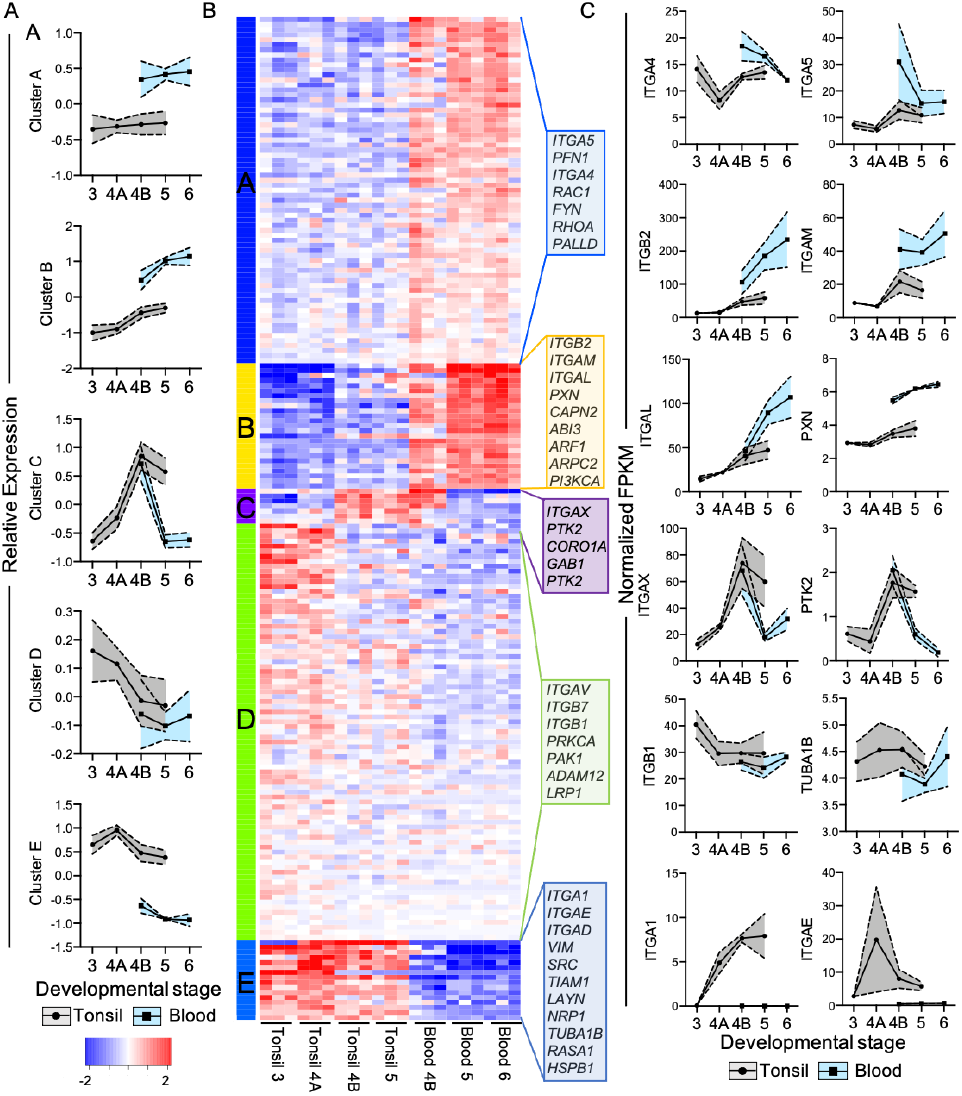
Human NK cell adhesome gene expression is determined by developmental stage and tissue residency. Adhesome gene expression data of tonsil and PB NK cells was subjected to K means clustering (K=5) and expression of genes within each cluster were averaged and plotted. A) Average expression of K means clusters of tonsils (grey) and PB (blue) NK cell subsets. Standard deviation is depicted by dashed lines. B) K means heatmap of adhesome gene expression of tonsil and PB NK cells subsets. C) Average gene expression of select representative adhesome genes from K means clusters. Ton, tonsil; PB, peripheral blood. Data derived from the means of 3 technical replicates pooled from 12 tonsil or peripheral blood donors.

### Integrin protein expression and activation reflects tissue specificity of NK cell developmental subsets

Our transcriptomic data suggested that there were significant differences in the expression of integrins between NK developmental subsets and sites of isolation. To define the cell surface expression of CD11a/CD18 [LFA-1], CD11b/CD18 [Mac-1], CD49d/CD29 [VLA-4] and CD49e/CD29 [VLA-5] integrins that were differentially expressed between these populations, we performed flow cytometric analyses of single cell suspensions from tonsil and PB donors (see Supp. Fig. 4 for representative gating strategy and Supp. Tables 3, 4 for antibodies and subsets). Our analysis of leukocyte specific integrins revealed that PB stage 3-6 and tonsil stage 3-5 NK cells from all donors were found to have cell surface expression of all β2 integrins tested (Fig. 3 histograms), with varying degrees of relative expression detectable by measuring mean fluorescence intensity (MFI). Expression of CD11a (integrin *α*L) was found to be significantly higher in NK cell stages 4B and 5 relative to stage 3 in tonsil, and PB stages 4-6 had significantly higher CD11a MFI relative to stage 3 NK cells (Fig. 3A). Interestingly, our transcriptomic data suggests that ITGAL (integrin *α*L) is more highly expressed in PB stage 5 but not stage 4 relative to tonsil (Fig. 2C), yet flow cytometric analysis revealed that CD11a was more highly expressed on the surface of both PB stage 4 and 5 NK cells relative to their counterparts in tonsil (Fig. 3A). Dissimilar to transcriptomic data, which shows increasing levels of integrin *α*M through PB NK cell maturation, analysis of PB NK cell surface expression of CD11b revealed that expression did not significantly vary between PB NK cell developmental subsets (Fig. 3B). Unlike in PB NK cell subsets, when we analyzed CD11b expression in tonsil NK cell subsets, we saw that tonsil NK cell subsets had a significant increase in the expression of CD11b between stage 5 and stage 3 NK cells (Fig. 3B). Expression of CD11c (integrin *α*X) did not significantly differ between tonsil NK cell stages (Fig. 3C), yet we observed a significant increase in CD11c between PB stages 3-4B (Fig. 3C), followed by a significant decrease in stage 5. Furthermore, CD103 (integrin *α*E) expression was consistent between developmental subsets in tonsil, while in PB, although this was not noted by transcriptomic analysis, CD103 was significantly upregulated between stages 3 and 6 as well as stages 4A and 6 (Fig. 3D). Lastly, we analyzed the expression of CD18 (integrin β2) and found that, in agreement with transcriptomic data, CD18 was significantly upregulated in more mature NK cell subsets stage 4B and 5 in tonsil and stage 5 and 6 in PB relative to stage 3 (Fig. 3E). Next, we sought to measure the cell surface expression of CD29 (integrin β1) and integrin β7 associated integrin subunits on phenotypically equivalent tonsil and PB NK cell subsets (Fig. 4). As suggested by our transcriptional data (Fig. 2C), PB NK cells do not significantly upregulate CD49a (integrin *α*1) (Fig. 4A). On the other hand, tonsil NK cell stages 4A-5 preferentially upregulated CD49a, which is associated with tissue residency, relative to tonsil stage 3 and PB counterparts (Fig. 4A). Both PB and tonsil NK cells did not variably express CD49d (integrin *α*4) through maturation, yet we found that PB NK cells stages 4 and 5 had increased expression of CD49d relative to their phenotypically equivalent counterpart in tonsil (Fig. 4B). CD49e (integrin *α*5), a β1 associated integrin receptor for fibronectin domains, is preferentially expressed by more immature stage 3 NK cells and is significantly downregulated through NK cell development in tonsil but to a lesser extent in PB (Fig. 4C). Moreover, both PB and tonsil NK cells downregulated CD29 as they matured from stage 3, albeit the downregulation of CD29 was greater in PB than in tonsil NK cells (Fig. 4D). These data, in parallel to our transcriptomic data, demonstrate that PB and tonsil NK cells differentially regulate cell surface expression of integrins both in a developmental and tissue residency dependent manner.

**Fig. 3.**
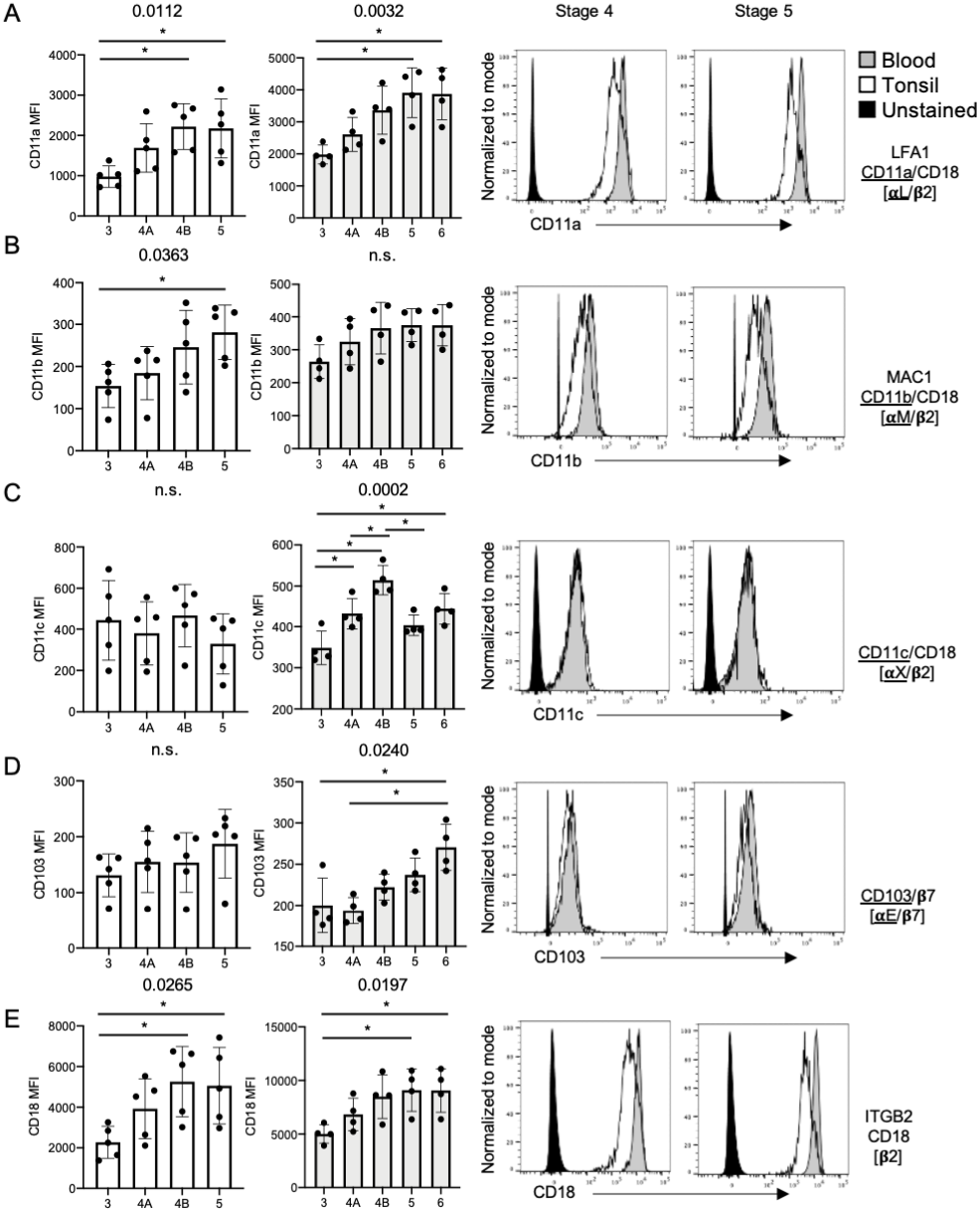
Leukocyte-specific integrin expression reflects the tissue specificity of human NK cell developmental subsets. Expression (MFI) of integrin subunits on NK cell developmental subsets was measured as described in Methods. Data shown are from 4-5 peripheral blood or tonsil healthy donors. A-E) MFI (left) and representative histograms (right) of leukocyte associated integrins CD11a, CD11b, CD11c, CD103, and CD18. Error bars indicate mean±SD. MFIs from each tonsil or PB NK cell subset were compared by ordinary one-way ANOVA with multiple comparisons. **P<0.05.

**Fig. 4.**
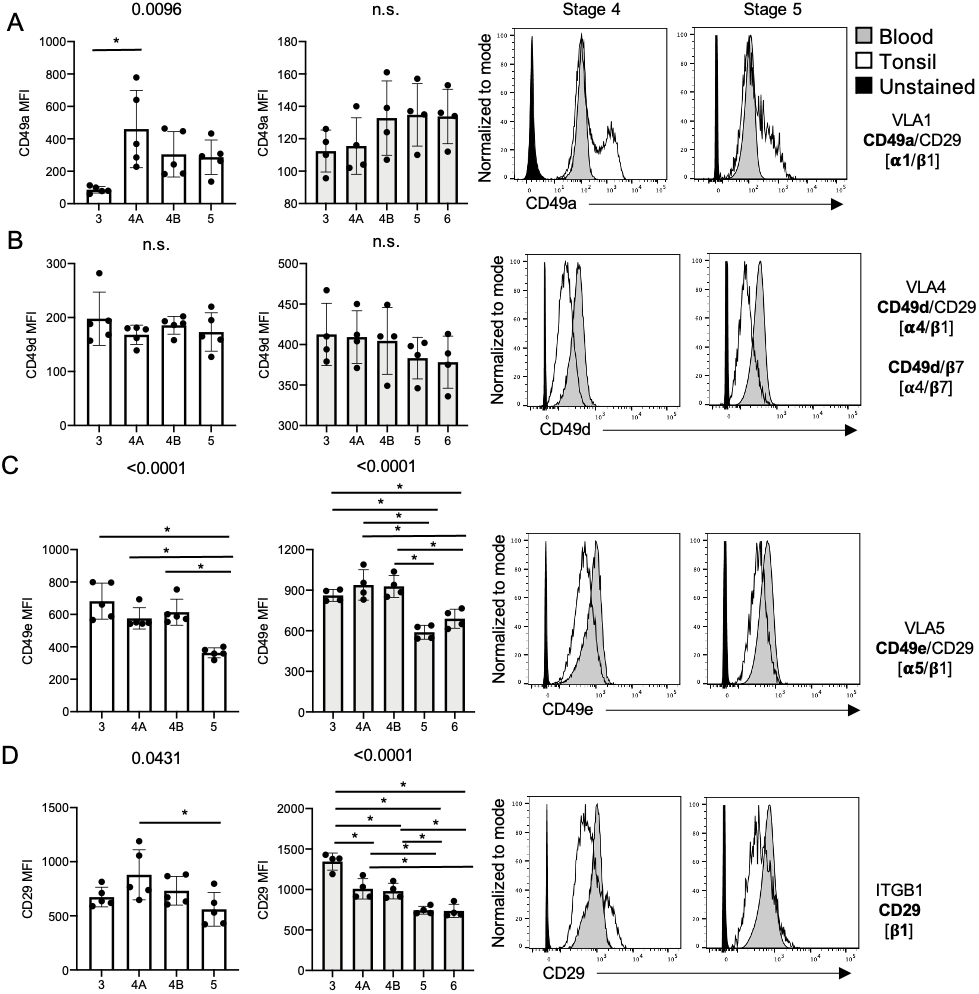
Integrin β1 and associated *α* subunit expression is dictated by NK cell developmental stage and tissue residency. Expression of selected β1 associated integrins, CD49a, CD49d, CD49e and CD29, from flow cytometry of dissociated tonsil mononuclear cells or peripheral blood mononuclear cells (center, right). Data shown are from 4-5 peripheral blood or tonsil healthy donors A-D) MFI and representative histograms of β1 associated integrins CD49a, CD49d, CD49e, and CD29. Error bars indicate mean±SD. MFIs from each condition were compared by ordinary one-way ANOVA with multiple comparisons. **P<0.05.

### Integrin conformation on NK cell developmental subsets

Based on the differential expression of integrins between NK cell subsets from peripheral blood and tonsil we wanted to understand how NK cells from these respective tissues regulate integrin activation, which has implicit effects on cell behavior, shape, and state (30–32). We included mAb24 and HUTS-4 antibodies for detection of open/extended conformation, or activated, CD11a-c/CD18 (LFA-1, MAC-1, integrin *α*X/β2), and CD49a-f/CD29 (VLA-1-6) respectively (39) (Fig. 5A, C). We found that tonsil NK cells had a greater frequency and MFI of cells with detectable open conformation of CD29 heterodimers when compared to analogous peripheral blood subsets, yet in PB we observe a significant increase in CD29 activation in stage 6 relative to more immature PB NK cells (Fig. 5A, B). We also observed an increased frequency of tonsil stage 4B NK cells with activated CD18 heterodimer compared to peripheral blood stage 4B NK cells (Fig. 5C, D). Taken together, our data suggest that NK cell subsets have differential expression and conformation of integrins on their cell surface, with unique integrin profiles that also reflect their tissue residency.

**Fig. 5.**
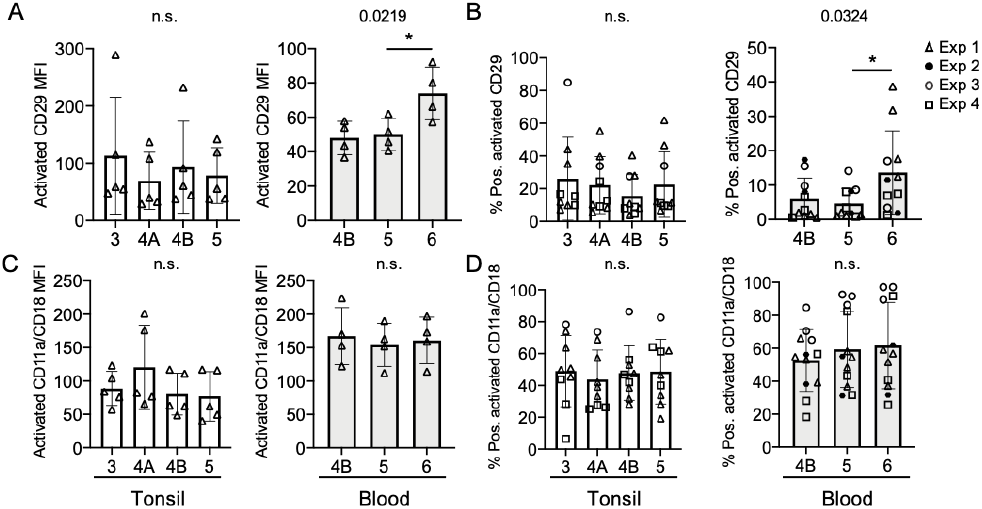
Integrin conformation is determined by tissue specificity and developmental stage. Flow cytometric analysis of dissociated tonsil mononuclear cells or peripheral blood mononuclear cells was used to measure expression (MFI) of activated integrin subunits. Data shown is of 5 tonsil and 4 peripheral blood healthy donors. Monoclonal antibodies that detect the open (activated) conformation of CD29 (CD49a-f/CD29) or CD18 (CD11a-c/CD18) were used to probe developmental subsets from tonsil or peripheral blood as indicated. Representative mean fluorescence intensity (A and C) or pooled frequency (B and D) of open conformation integrins on NK cells were quantified relative to unstained samples. Representative MFI data shown are from 4 peripheral blood and 5 tonsil donors analyzed on the same day. 11-12 PB and 9 tonsil donors run on separate days as indicated by the shape of the data point. Normally distributed frequencies of cells with activated CD29 or CD18 were compared by ordinary one-way ANOVA with multiple comparisons, while non-parametric data was analyzed by Kruskal-Wallis Test with multiple comparisons. MFIs from each condition were compared by ordinary one-way ANOVA with multiple comparisons **P<0.05.

### Expression of integrins on in vitro derived NK cells resembles in vivo patterns

In vitro differentiation of NK cells from CD34^+^ precursors has been well described as a method of studying human NK cell development (40, 41). While innate lymphocytes undergoing maturation in such systems are thought to progress through stages of differentiation similarly to those in situ (9, 12, 42, 43), the use of xenogenic stromal cells and exogenous cytokines in such systems make it difficult to define the role of the microenvironment in this process. Given the differences between analogous subsets of NK cells isolated from different tissues, we sought to define the expression of integrins on NK cells generated from in vitro differentiation. Primary human CD34^+^ cells from peripheral blood were isolated (Supp. Fig. 5) and cultured with EL08.1D2 or OP9 stromal cell lines in the presence of cytokines [FLT3L, SCF, IL-3, IL-7, IL-15] (40–43). Following 4 weeks of in vitro NK differentiation, stage 4 and 5 NK cells represented a significant proportion of the CD45^+^ population (Supp. Fig. 6, 7). We performed flow cytometry with our integrin panel described above after 4 weeks of differentiation on EL08.1D2 or OP9 stromal cells. At this timepoint the frequency of stage 1 and 2 NK cells significantly decreases relative to mature stages 3-5 which represent over 50% of the culture, thus only stage 3-5 NK cells were analyzed (Supp. Fig. 7). As with NK cells from tonsil and PB, CD18 (integrin β2) expression increased with maturation at stages 4 and 5, while CD29 (integrin β1) expression decreased (Fig. 6A). In addition, the MFIs of activated conformations followed a similar trend, showing an increase in open conformation CD18 heterodimers and a decrease in open conformation CD29 heterodimers through development (Fig. 6B, C). Notably, we did not observe a significant difference in integrin expression or conformational activation between in vitro derived NK cells that were generated by culture with OP9 or EL08.1D2 feeder cells. Finally, given the changes in integrin adhesome expression between developmental subsets, we sought to define the density of cellular actin in NK cells generated in vitro. We found that as NK cells matured, they exhibited a significant increase in density of F-actin content following cell permeabilization as measured by intracellular detection of phalloidin by flow cytometry (Fig. 6D). Together our results indicate that in vitro derived NK cells have conserved patterns of integrin expression when compared to ex vivo NK cells and exhibit differential F-actin content in a stage-specific manner.

**Fig. 6.**
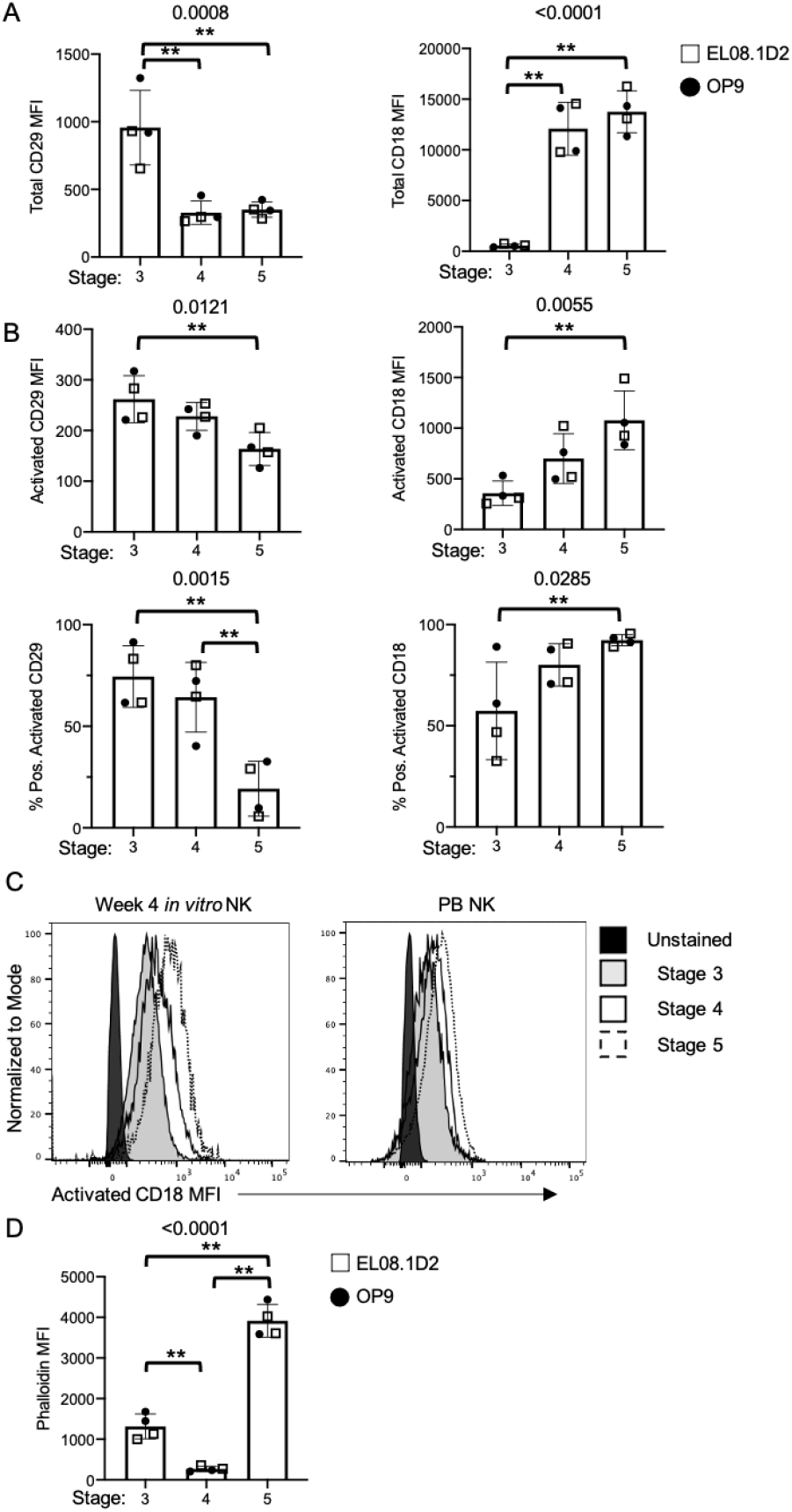
In vitro derived human NK cell developmental intermediates exhibit developmental stage-specific expression of integrins reflective of ex vivo subsets. NK cells were differentiated in vitro with co-culture on EL08.1D2 or OP9 stromal cells and their phenotype was analyzed weekly by flow cytometry. A) Expression (MFI) of CD18 and CD29 on in vitro-derived NK cells cultured with EL08.1D2 (open squares) or OP9 (filled circles) stromal cells. n=2 biological and technical repeats per stromal cell condition. B) MFI (top) or percent positive NK cells (bottom) of open conformation CD18 or CD29 on in vitro derived NK cell subsets measured by flow cytometry. n=2 biological and technical in vitro NK replicates. Statistical significance was tested by ordinary one-way ANOVA with multiple comparisons. Number above graph indicates *P* value with significant individual comparisons marked by **P<0.05. C) Representative histogram of activated CD18 heterodimer expression (MFI) on stage 3 (solid line, black), stage 4 (solid line, white) and stage 5 (dashed line, white) of in vitro derived (left) or peripheral blood (right) NK cells. D) Intensity of phalloidin detected by intracellular staining of in vitro derived NK cells at week 4 measured by flow cytometry. **P<0.05 by ordinary one-way ANOVA with multiple comparisons. Error bars indicate mean±SD.

### Primary NK cell subsets have distinct cortical actin densities

Given that flow cytometric analysis of in vitro derived NK cells unexpectedly revealed increased intensity of actin in mature NK cells, we sought to investigate the nature of cortical actin in ex vivo NK cells. Components of the actin cortex include actin, myosin, and regulators of actin polymerization and turnover, including the Arp2/3 complex and formins. Pools of actin monomers provide substrate for the generation and maintenance of the filamentous (F-) actin cortex, as well as the leading edge and immune synapse. Non-muscle mammalian cells contain two structurally similar isoforms of actin monomers, g-actin and b-actin, encoded by the *ACTG1* and *ACTB* genes respectively (44, 45). Given the ubiquitous nature of actin within all cells, it was unsurprising that we found that while there was a trend towards increased *ACTB* expression in more terminally mature NK cell subsets, there was no significant difference in expression of *ACTB* or *ACTG1* between the isolated subsets of human NK cells in our bulk RNA-Seq dataset (Fig. 7A). To more deeply characterize the nature of filamentous actin in peripheral blood and tonsil NK cells we performed intracellular detection of phalloidin by flow cytometry as we had previously done for in vitro-derived NK cells (Fig. 6). Strikingly, we found that there was a clear demarcation in actin content, with stage 5 NK cells having higher intensity of phalloidin detection than stage 4 cells (Fig. 7B). More extensive analysis of both tonsil and peripheral blood NK cell subsets showed that actin content was higher in NK cells from stages 5 and 6 in peripheral blood, whereas tonsil NK cell subsets had higher density of phalloidin staining in stage 6 NK cells (Fig. 7C). Given the differences we found by flow cytometry we sought to directly visualize and measure the cortical actin network in freshly isolated peripheral blood NK cells. We performed super-resolution structured illumination microscopy of actin by phalloidin detection in freshly isolated stage 4 and stage 5 cells from peripheral blood. Imaging confirmed our flow cytometry results and we observed a significantly greater density of cortical actin in more mature cells (Fig. 7D). As predicted, CD56 intensity measured by integrated density (MFI*area) was significantly greater in stage 4 CD56^bright^ NK cells than stage 5 CD56^dim^ NK cells (Fig. 7E). Similar measurements of actin further validated our observations as we found that integrated density of actin was significantly greater in stage 5 NK cells (Fig. 7F). While such measurements could be reflective of differences in cell size, we found that CD56^dim^ NK cells did not have a significantly greater volume than CD56^bright^ cells (Fig. 7G). Therefore, stage 5 NK cells have greater actin density relative to stage 4 cells and this is independent of cell size or activation. Finally, we sought underlying differences in gene expression that could drive the differences in actin density that we observed between NK cell developmental subsets. Further analysis of our RNA-Seq datasets identified key actin nucleation promoting factors and cytoskeletal regulators that were differentially expressed, including *ARPC2* (the ARPC2 subunit of the Arp2/3 complex) and *PFN1* (profilin) (Fig. 7H). Therefore, while there are a number of adhesome related genes that function in adhesion and migration that have different patterns of differential expression between tissue and peripheral blood, actin nucleating pathways seem to be specifically upregulated in terminally mature NK cells in peripheral blood. This is reflected by a greater density of cortical actin that is detectable in the absence of cellular activation and not linked to expression of actin monomers *ACTB* and *ACTG1*.

**Fig. 7.**
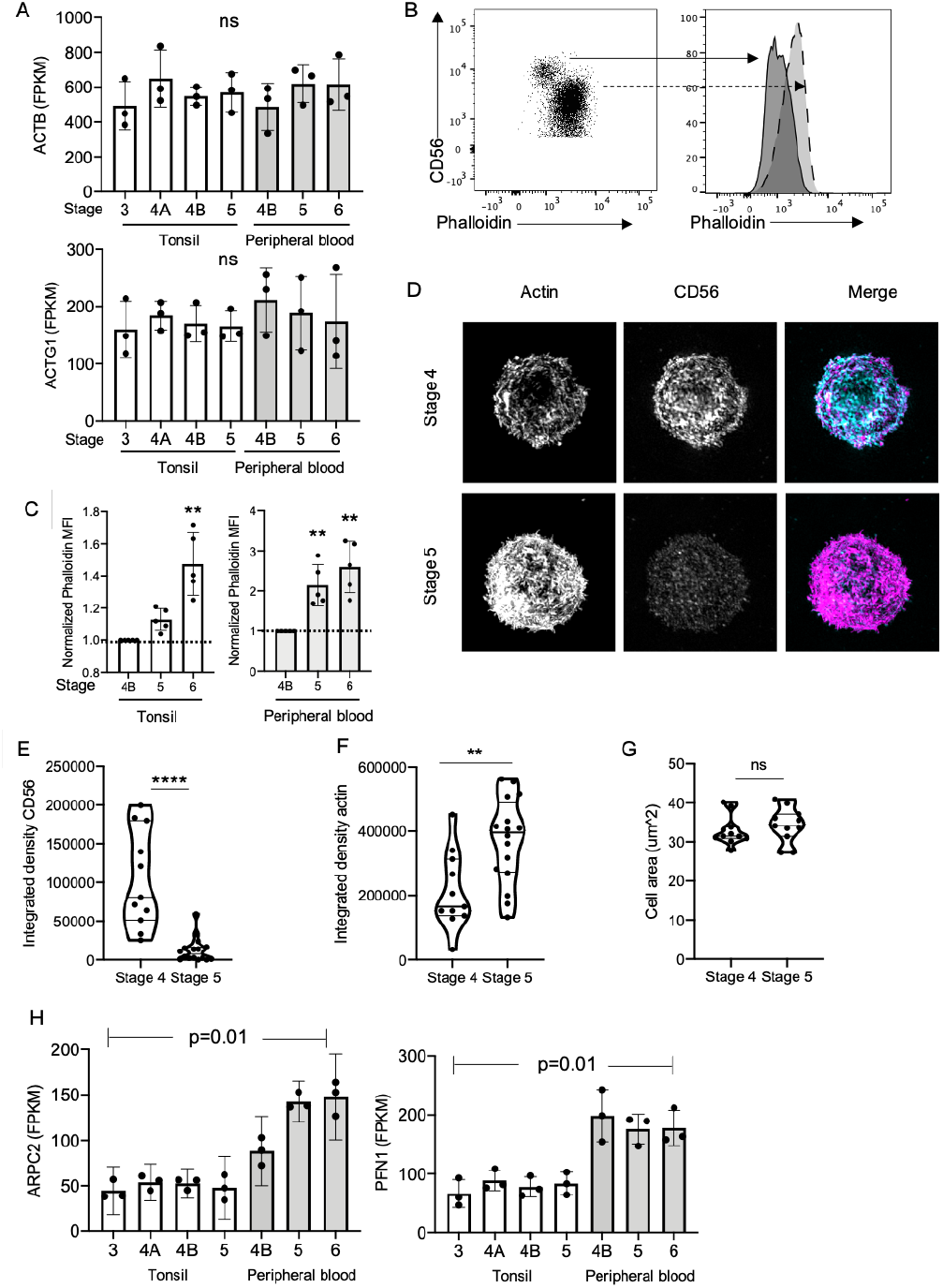
Terminally mature primary NK developmental subsets have increased density of cortical actin. Primary NK cells were isolated from tonsil and subsets were isolated as described in Methods. A) *ACTB* (top) or *ACTG1* (bottom) gene expression (FPKM) from RNA-Seq data of tonsil and peripheral blood NK cell developmental intermediates. n=3 biological replicates, mean±SD. ns, not significant by Kruskal-Wallis test. B) Representative flow cytometric analyses of phalloidin content in peripheral blood NK cells. C) Quantification of actin intensity in NK cell developmental subsets from tonsil or peripheral blood. n=5 biological replicates, mean±SD. **P<0.05 by Kruskal-Wallis Test with multiple comparisons. D) Structured illumination microscopy images of primary NK cells freshly isolated, fixed, permeabilized and immunostained with phalloidin (magenta) and anti-CD56 antibody (cyan). E) CD56 integrated density. n=11 (CD56bright), 16 (CD56dim); ****P<0.0001 by Mann-Whitney test. F) Actin integrated density. n = 11 (CD56bright), 16 (CD56dim); ***P<0.001 by Mann-Whitney test. G) Cell volume of stage 4 and stage 5 cells. n=10 per condition; ns, not significant by Mann-Whitney test. H) Normalized estimated gene expression (FPKM) of selected genes *ARPC2* and *PFN1* related to actin remodeling and nucleation. P = 0.01 by Kruskal-Wallis test.

## Discussion

Our data suggest that NK cells receive both intrinsic and extrinsic cues which dictate integrin adhesome expression. Tonsil NK cells are representative of secondary lymphoid tissue resident NK cells (8, 10), suggesting that the expression of adhesome components involved in migration and ECM interactions, such as collagen-specific integrins, are preferentially expressed in these tissues relative to peripheral blood. Conversely, peripheral blood NK cells upregulate integrin adhesome components that promote cellular integrity in an environment with constant shear flow and enable cells to bind endothelial cells lining blood vessels during extravasation (39). Previous studies have found that *ITGA1, ITGAE* and *IT-GAD*, which are all upregulated in tonsil NK cell populations relative to peripheral blood NK cells, are associated with lymphocyte tissue residency and migration (5, 21, 46, 47). ITGAE (CD103) is upregulated in tissue resident memory CD8^+^ T cells from lung and spleen, as well as in CD56^bright^CD16^−^ NK cells from tissue sites such as lung, endometrium, nasal mucosa and intestine, thus providing support for our observations that the adhesome expression of tonsil NK cells includes that of previously characterized tissue resident lymphocytes (5, 23, 46, 47). While less well characterized than integrin *α*E, integrin *α*D is expressed on human NK cells and is associated with increased adhesion to ECM components, inside-out signaling-dependent cytokine secretion, and homing to inflammatory sites (48). Our ability to distinguish stage 4A and 4B NK cells and define the up-regulation of β2 integrins, such as *ITGAL* (CD11a), *ITGAM* (CD11b), and *ITGB2* (CD18), as stage 4A NK cells mature into stage 4B NK cells in tonsil allowed us to further define differences in adhesome profiles that occur at this stage of maturation associated with commitment to NK cell maturation (13). Based on these observations, and the higher expression of cytoskeletal signaling and regulator genes including focal adhesion kinase (*PTK2*), calpain 2 (*CAPN2*), and *TIAM1*, we show that tonsil NK cells have a unique adhesome profile that is defined by higher expression of both integrins and the cytoskeletal machinery that mediates their interactions with tissue.

Our data demonstrate that the largest changes in adhesome gene expression in both peripheral blood and tonsil occur early in NK cell development, and we found many adhesome genes differentially expressed between peripheral blood stages 4B and 5, while none were differentially expressed between tonsil stages 4B and 5. This suggests that there are still unknown differences between representative stage 4B and 5 NK subsets in tonsil and peripheral blood that go beyond the differences in expression of NK markers commonly used to discriminate developmental subsets between tissues. Still, our observations are in line with previous studies showing that tonsil- and lymph node-resident stage 5 NK cells resemble a more immature-like stage 4B NK cell, with increased expression of CD56 and decreased expression of cytotoxic function-related machinery, whereas stage 4B and stage 5 NK cell subsets in peripheral blood are unique both transcriptionally and phenotypically (4, 5).

When considering these differences between NK cells isolated from different sites, we sought to better define how in vitro derived cells align with primary cells. In vitro studies have demonstrated a functional requirement for integrins, specifically VLA-4 (*α*4β1), in facilitating T cell precursor interactions with OP9-DL1 stroma and priming double negative T cells to receive Notch signaling (49). When we consider integrin β1 and integrin β2 expression, we observed that, similar to primary human NK cells, in vitro-derived NK cells have changes in integrin expression between stages 3 and 4 of development. Specifically, NK cells transition from having relatively high integrin β1 in early stages of in vitro NK cell development to having low integrin β1 and high integrin β2 expression during stages 4-5. The transition between high expression of integrin β1 to integrin β2 suggests that the upregulation of integrin β2 is a key feature of NK maturation in vitro and occurs in concert with the acquisition of CD94 and downregulation of CD117 (c-kit). The similarities in the patterns of changes of integrin β1 and integrin β2 density that we observe between in situ- and in vitro-derived NK cells suggest that relative density of integrin expression may be an intrinsically programmed feature of NK cell maturation. In contrast to the expression of total integrin subunits, we found that the relative frequencies of cells with activated conformation of CD18 and CD29 heterodimers differed between in vitro and in situ-derived cells. Measuring the amount of activated integrin on the surface of NK cells allowed us to understand differences in the regulation of these subunits between tissue sites and demonstrated that integrin activation is dependent on both the microenvironment and developmental stage of NK cells. Factors that could contribute to the observed differences in integrin activation between in vitro and in situ-derived cells include differences in ECM components and the exogenous use of cytokines, particularly IL-15 (50).

Our finding that NK cells at later stages of development, namely stage 5 in peripheral blood and stages 4 and 5 in vitro, had increased density of cortical actin was surprising. While we did see a slight increase in the expression of the actin monomer *ACTB*, the difference in cortical actin density as measured by both flow cytometry and super-resolution microscopy was orders of magnitude greater in stage 5 (CD56^dim^) NK cells than stage 4 cells (CD56^bright^). Further, it should be noted that we did not activate the cells with integrin or activating receptor ligation. As such, the differences we observed were that of the cortical actin meshwork, not activation-induced actin remodeling. In contrast with other cell surface receptors such as L-selectin and CD94 that appear to have intermediate expression as cells progress from stage 4 to stage 5 (11, 51), we found a sharp demarcation in phalloidin intensity between CD56^bright^ and CD56^dim^ cells. Shear flow, such as that found in circulation, induces rapid lymphocyte morphological changes prior to tissue extravasation, suggesting that the CD56^dim^ NK cells found primarily in circulating peripheral blood may have increased cortical density in response to shear stresses or actin polymerization induced by tethering or transendothelial migration. However, given that the CD56^bright^ (stage 4) population isolated from peripheral blood had uniformly lower phalloidin staining, it seems unlikely that this is the case unless CD56^dim^ NK cells have uniquely undergone transendothelial migration. The increased expression of actin nucleating proteins, including actin-related protein 2/3 complex subunit 2 and profilin, in stage 5 peripheral blood cells suggests that the higher expression of these correlates with this higher actin density, and ARPC2 is a known regulator of cortical actin thickness (52). More detailed investigations into actin dynamics and architecture in primary NK cells will be necessary to link the phenotypic differences that we observed with functional outcomes.

While here we have been guided by the consensus integrin adhesome, what remains to be defined is the spatial information regarding the formation of integrin adhesion complexes and signaling islands. Studies of the integrin adhesome are generated by proteomic data and further probed by high- and super-resolution microscopy that provides critical spatial information about how such complexes are formed and function (28–30, 32, 53, 54). An additional caveat of our approach is that both flow cytometry and gene expression data consider only discrete integrin subunits, yet their function and ligand specificity must be considered in the context of their obligate heterodimeric structure. However, by defining the expression of adhesome components at the gene expression level, and validating some of these by protein expression, we lay the foundation for future studies that will better define the role of integrins through multi-scale approaches.

## Methods

### Primary NK cell isolation from peripheral blood and tonsil samples

All human tissues used in the RNA-sequencing studies were collected under a protocol approved by The Ohio State University Institutional Review Board. Human pediatric tonsils were obtained fresh through the Cooperative Human Tissue Network (CHTN) from Nation-wide Children’s Hospital (Columbus, OH), and peripheral blood was obtained through the American Red Cross as previously described (13, 33). Single cell suspensions were enriched for NK lineage cells using a bivalent antibody RosetteSep (StemCell Technologies)-based method (55), and then the resultant enriched NK cell fractions were labeled with antibodies (see Supp. Table 3 for a complete list of flow antibodies) and finally sorted to purity using a BD FACS Aria II cell sorter. Purities were validated post-sort and all samples had purity greater than 99%. Human tonsil NK cell subsets were defined and sorted as follows: stage 3 (Lin^−^CD117^+^CD94^−^NKp80^−^CD16^−^), stage 4a (Lin^−^CD94^+^NKp80^−^CD16^−^), stage 4b (Lin^−^CD94^+^NKp80^+^CD16^−^), stage 5 (Lin^−^NKp80^+^CD16^+^); human blood NK cell subsets were defined and sorted as follows: stage 4b (Lin^−^NKp80^+^CD16^−^CD57^−^), stage 5 (Lin^−^NKp80^+^CD16^+^CD57^−^), stage 6 (Lin^−^NKp80^+^CD16^+^CD57^+^). For these sorting experiments, Lin = CD3, CD14, CD19, CD20, CD34.

For flow cytometry, whole blood was obtained by venipuncture from healthy donors or as discarded apheresis product from patients undergoing routine red blood cell exchange at Columbia University Medical Center. Alternatively, we acquired RBC-low Leukocyte-enriched Buffy Coats from the New York Blood Bank as an alternative source of primary NK cells for flow cytometric analysis. Primary NK cell subsets from blood were enriched with NK cell RosetteSep (StemCell Technologies). Blood was layered onto Ficoll-Paque density gradient followed by centrifugation at 2,000 rpm for 20 mins (no brake). NK cells were collected from the density gradient interface and washed with PBS by centrifugation at 1,200 rpm for 7 mins. NK cells were resuspended in PBS 10% FCS and counted, then either resuspended in PBS for flow cytometry or cryopreserved in fetal calf serum with 10% DMSO at a concentration of 1 - 2.5 × 10^6^ cells per ml.

Tonsil samples for dissociation and flow cytometry analysis were acquired from routine tonsillectomies performed on pediatric patients at Columbia University Irving Medical Center. Tissue samples were placed in a sterile dish with PBS and manually dissociated by mincing into a cell suspension. The cell suspension was then passed through a 40 μm filter to obtain a single cell suspension and washed with PBS by centrifugation at 1,200 rpm for 7 mins. Cells were either resuspended in PBS for flow cytometry or FCS 10% DMSO freezing media at a concentration of 5-10 × 10^6^ cells and cryopreserved prior to use.

CD34^+^ precursors for in vitro experiments were isolated from whole blood obtained by venipuncture from two healthy donors. Mononuclear blood cells were isolated from donors were incubated with anti-CD34 antibody (Supp. Table 3) prior to cell sorting. CD34^+^ cells were isolated by FACS sorting on a BD Aria II cytometer with an 85 μm nozzle at 45 p.s.i. Sorted cells were confirmed to be >90% CD34^+^ and were cultured directly after isolation on previously irradiated EL08.1D2 or OP9 cells as described below. Primary cells, apheresis and tonsils were obtained in accordance with the Declaration of Helsinki with the written and informed consent of all participants under the guidance of the Institutional Review Boards of Ohio State University and Columbia University.

### Peripheral blood and tonsil NK cell bulk RNA sequencing and analysis

Freshly sorted blood and tonsil NK cells were pelleted, and total RNA was isolated using the Qiagen RNeasy Mini Kit (Qiagen, Active Motif). Directional poly-A RNA sequencing libraries were prepared and sequenced as 42-bp paired-end reads on an Illumina NextSeq 500 instrument (Illumina) to a depth of 33.2 – 48.0 x 10^6^ read pairs (Active Motif). Alignment to human genome (hg19 build) was done using TopHat. Transcriptome assembly and analysis was performed using Cufflinks and expression was reported as FPKM.

Fragments per kilobase of transcript per million mapped read (FPKM) gene expression data were obtained from prefiltered and normalized bulk RNA sequencing raw data and imported to iDEP 0.9 (56). FPKM bulk RNA seq data (Supp. Table 5) was processed using the source code available on iDEP 0.9 (56) following the recommended parameters. Pathway analysis was performed on iDEP 0.9 transformed RNA seq data using PGSEA ranking and KEGG pathways (56, 57), filtering on pathways with 15-2000 genes and FDR cutoff of 0.2. Prism 8.0 (GraphPad Software) was used to visualize data.

### In vitro NK cell differentiation

EL08.1D2 cells were a gift from Dr. Jeffrey Miller (University of Minnesota) and were cultured as previously described (58) in culture flasks pre-treated with 0.1% gelatin. Cells were maintained at 32°C in 40.5 α-MEM (Life Technologies), 50% Myelocult M5300 (StemCell Technologies), 7.5 heat-inactivated fetal calf serum (Atlanta Biologicals) with β-mercaptoethanol (10^*−*5^ M), Glutamax (Life Technologies, 2 mM), penicillin/streptomycin (Life Technologies, 100 U ml^*−*1^), and hydrocortisone (Sigma, 10^*−*6^ M), supplemented with 20% conditioned media. OP9 cells (ATCC) were cultured in non-gelatinized culture flasks at 37°C in alpha minimal essential media with 20% heat inactivated FBS and 1% Penicillin/Streptomycin (Life Technologies, 100 U ml^*−*1^). Prior to in vitro NK cell differentiation, 10^4^ EL08.1D2 cells were seeded into 96-well flat bottom plates pre-coated with 0.1% gelatin, while OP9 cells were similarly seeded into non-gelatinized 96-well flat bottom plates. Cells were grown to confluence then subjected to mitotic inactivation by irradiation at 30 Gy. Following FACS sorting, CD34^+^ cells were cultured in NK cell differentiation media containing Ham F12 media plus DMEM (1:2) with 20% human AB-serum, ethanolamine (50 μM), ascorbic acid (20 mg ml^*−*1^), sodium selenite (5 μg ml^*−*1^), β-mercaptoethanol (24 μM) and penicillin/streptomycin (100 U ml^*−*1^) in the presence of IL-15 (5 ng ml^*−*1^), IL-3 (5 ng ml^*−*1^), IL-7 (20 ng ml^*−*1^), Stem Cell Factor (20 ng ml^*−*1^), and Flt3L (10 ng ml^*−*1^) (all cytokines from Peprotech). CD34^+^ cells were seeded onto irradiated EL08.1D2 or OP9 at a density of 2 × 10^3^ cells per well and incubated at 37°C with weekly half media exchanges.

### Primary and in vitro derived NK cell flow cytometric analysis

Flow cytometry to quantify integrin expression was performed using antibodies as described in Supplemental Table 3 and Fig. 5. Cryopreserved primary NK cells were thawed and resuspended in RPMI 10% FCS then immunostained at the concentrations indicated. For intracellular phalloidin staining cells were first incubated with antibodies for surface receptors, fixed and permeabilized using CytoFix/Cytoperm (BD Biosciences), then incubated with directly conjugated phalloidin. Data were acquired on a Bio-Rad ZE5 Cell Analyzer then exported to FlowJo 10 (BD Biosciences) for analysis. Integrin subunit MFI of primary tonsil and peripheral blood NK cell subsets were used to directly compare integrin expression of populations of cells collected on the same day. For flow cytometry of in vitro derived NK cells, cells were isolated at weekly time points and immunostained (Supplemental Table 3). NK developmental subsets were identified and analyzed using the gating strategy described in supplemental figures 4, 5, 6 and 7. Data were plotted and statistical analysis was performed using Prism 8.0 (GraphPad Software).

### Microscopy and image analysis

For structured illumination microscopy primary NK cells were enriched from peripheral blood of healthy human donors using RosetteSep (Stemcell Technologies). Freshly isolated cells were pre-incubated with anti-CD56 Alexa Fluor 647 (clone HCD56, Biolegend, 1:100) for 20 minutes in a conical tube then incubated on #1.5 coverslips that had been pre-coated with poly-L-lysine for an additional 20 minutes at 37°C 5% CO2. Following incubation, cells were fixed on the coverslip with BD CytoFix/CytoPerm for ten minutes. Cells were gently washed with PBS 2% FCS containing 0.1% saponin then incubated for 30 minutes with phalloidin Alexa Fluor 568 (Thermo, 1:100). Coverslips were gently washed again then mounted with Prolong Gold (Thermo). Imaging was performed on a GE DeltaVision OMX SE in 3D-SIM mode through a 60X 1.42 NA APO objective. 3D images were captured at 125 nm steps and the pixel size was 79 nm. Images were reconstructed using 3 orientations and 5 phase shifts with a Wiener filter constant of 0.001. Acquisition and reconstruction were performed with GE SoftWoRx software and images were exported to Fiji (59) for further analysis. Calculation of integrated density was performed by applying a uniform default auto threshold in Fiji then measuring area, intensity and integrated density (area*MFI) for CD56 and actin. Data were graphed and statistics were calculated in Prism 8.0 (GraphPad Software).

### Statistics

To test the normal distribution of flow cytometric data we utilized Shapiro-Wilk Test, based on our sample size, with a P value cut off of 0.05. Our MFI and cell frequency data fell under a normal distribution. Thus, we used an ordinary one-way ANOVA with multiple comparisons to compare the MFIs or cell frequency of our NK cell developmental subsets (Fig 3, 4, 5, 6). For our non-parametric data, as determined by a Shapiro-Wilk test *P* value less than 0.05, we employed a Kruskal-Wallis test with multiple comparisons to compare the mean frequencies of CD29 activation of our NK cell developmental subsets from tonsil. We also used a Kruskal-Wallis test with multiple comparisons test to compare the normalized means of all developmental stages and specifically stage 5 and 6 phalloidin intensity to that of stage 4B. Phalloidin intensity of PB and tonsil NK cells was normalized to the MFI of stage 4B cells within the respective sample. Our P value cutoff for significance of all statistical tests comparing the means of samples was 0.05. Mann-Whitney tests were used to compare unpaired data with a non-normal distribution identified by Shapiro-Wilkes test. All statistical testing was performed in Prism (GraphPad Software).

## ACKNOWLEDGEMENTS

The authors would like to thank Michael Kissner for technical support and acknowledge the use of shared resources of the Columbia Stem Cell Initiative flow cytometry core and the Herbert Irving Comprehensive Cancer Center Radiation Research core. The authors wish to thank Evelyn Hernandez and Carlos Aguilar Breton for their assistance with procuring and processing tonsil samples. This work was supported by grants from the National Institutes of Health (R01AI137073 to E.M.M., and R01CA199447, R01CA208353 to A.G.F.). Research reported in this publication was also supported by The Ohio State University Comprehensive Cancer Center and the National Institutes of Health (P30-CA016058). The authors thank the Cooperative Human Tissue Network of Nationwide Children’s Hospital (Columbus, OH) for human pediatric tonsil samples. We also thank Ricardo Henriques for making his bioRxiv template freely accessible.

---

**Supplemental Figure 1:**
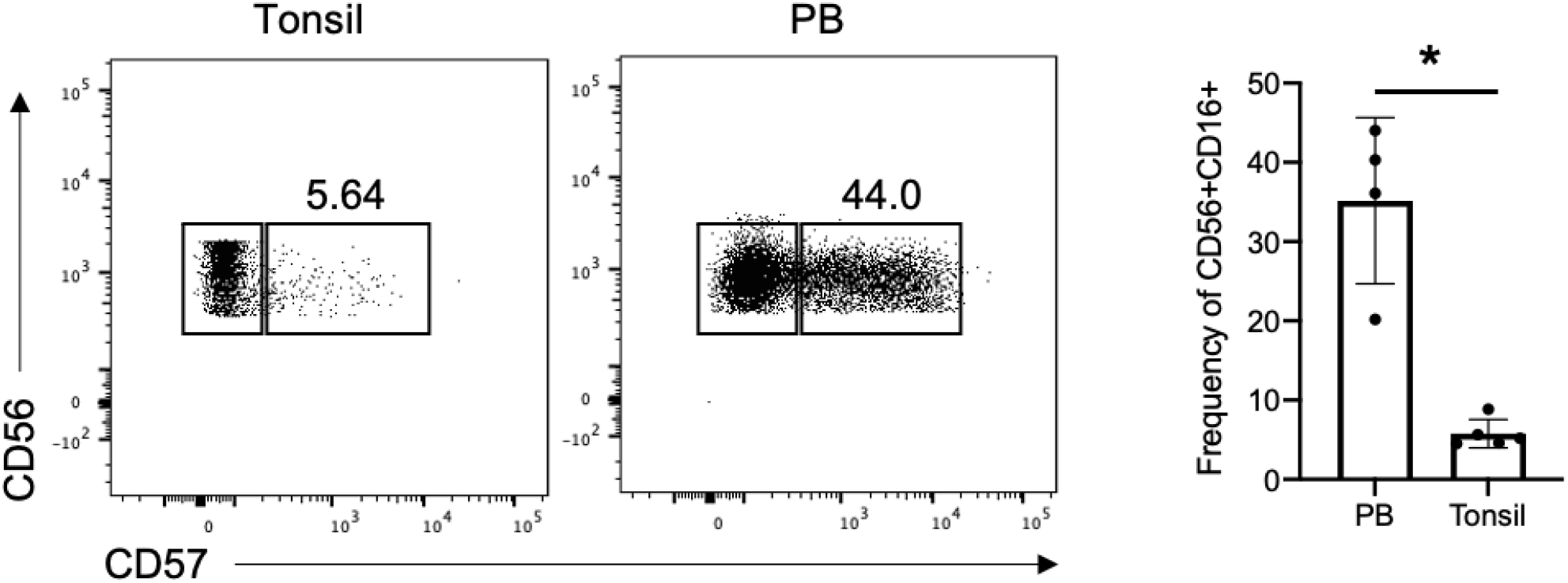
Frequency of CD57^+^CD16^+^CD56^dim^ stage 6 NK cells and other developmental subsets in tonsil and PB. **A)** Representative dot plots of stage 5 and 6 cell counts in PB and tonsil samples, gated on CD45^+^CD3“CD18”CD14’CD117“CD94^+/l0W^CD56^dim^CD16^+^. **B)** Frequency of NK stage 6 subsets within CD56^dim^CD16^+^ NK cells were calculated from 5 Tonsil and 4 PB donors. Means were compared using Mann-Whitney test P value=0.0159.

**Supplemental Figure 2:**
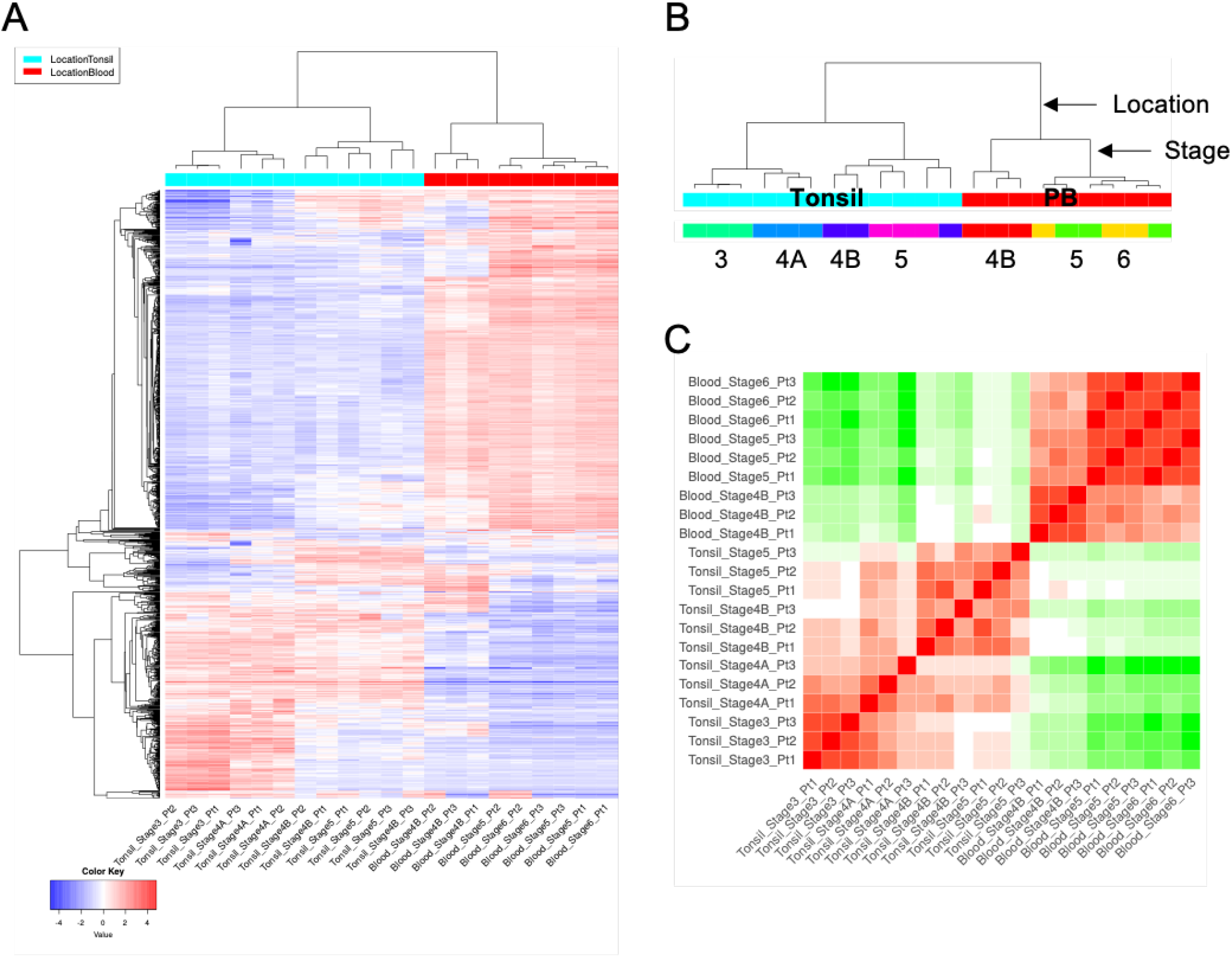
NK cell gene expression is dependent on developmental stage and tissue residency. **A)** Hierarchical heatmap of top 2,000 variably expressed genes of peripheral blood and tonsil NK cell subsets, not filtered on adhesome genes. **B)** Schematic of hierarchical clustering of peripheral blood and tonsil NK cell subsets based on total gene expression. **C)** Correlation matrix of peripheral blood and tonsil NK cells based on top 2,000 variably expressed genes ranked by standard deviation.

**Supplemental Figure 3:**
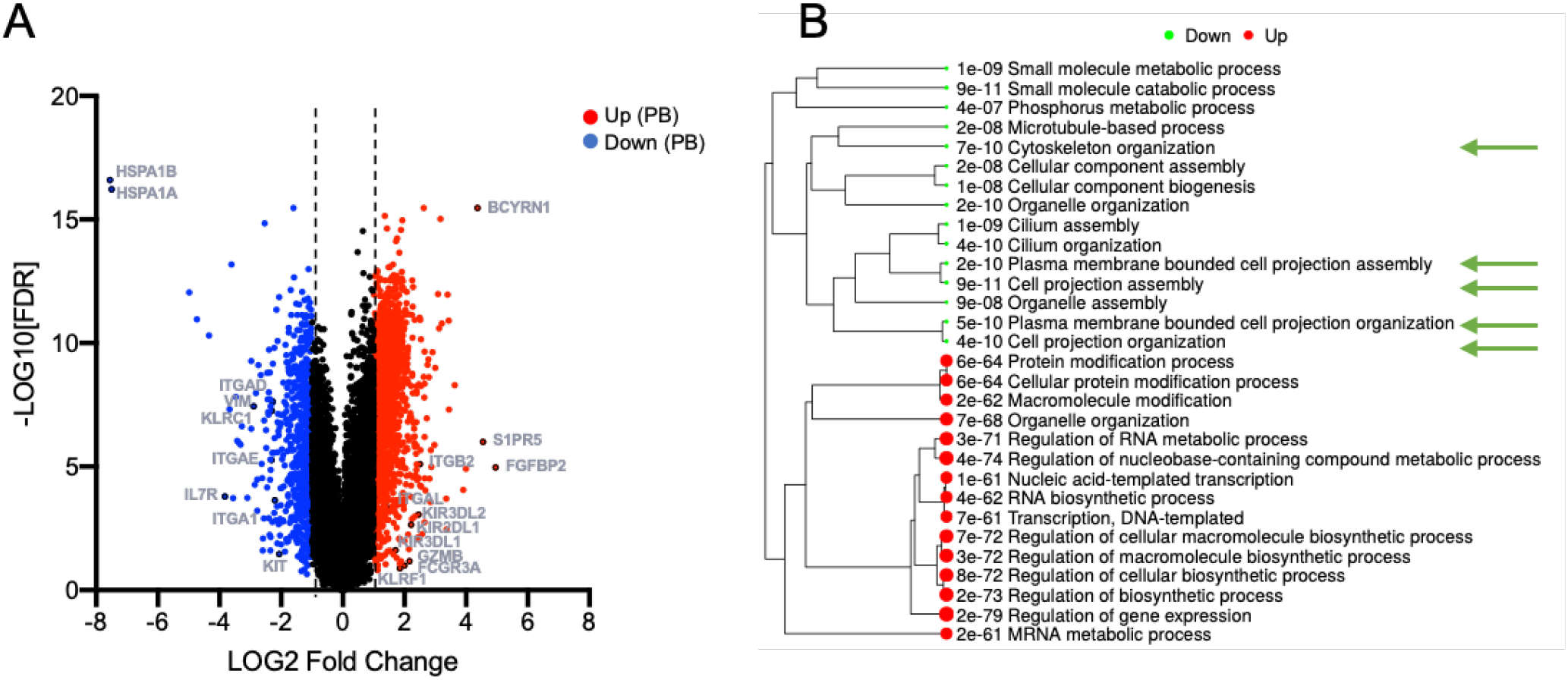
Comparison of peripheral blood and tonsil NK cell gene expression reveals that adhesome related genes are variably expressed between tissue sites. **A)** Differential gene expression analysis of peripheral blood and tonsil NK cells highlighting genes with ≤2-fold cutoff and FDR ≥0.1 (red = up in peripheral blood, blue = down in peripheral blood) **B)** GO Biological Process Pathway analysis of top differentially expressed genes in peripheral blood relative to tonsil NK cells. Green arrows indicate pathways related to adhesome genes.

**Supplemental Figure 4:**
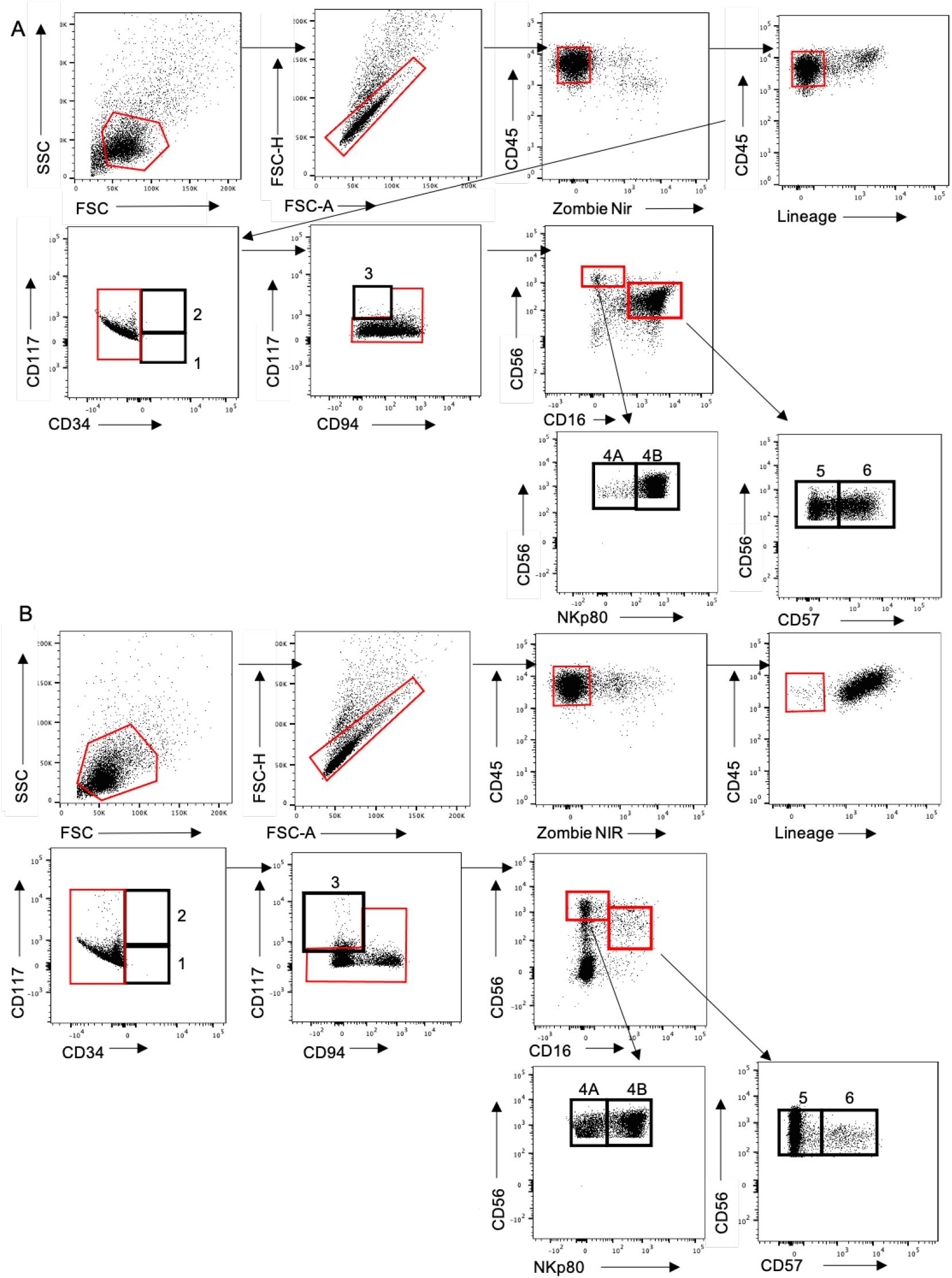
Representative flow cytometric gating strategy for analyzing peripheral blood and tonsil NK cell subsets. **A)** peripheral blood NK cells. **B)** Tonsil NK cell subsets. Numbers indicated NKcell development stages g isolated with each gate.

**Supplemental Figure 5:**
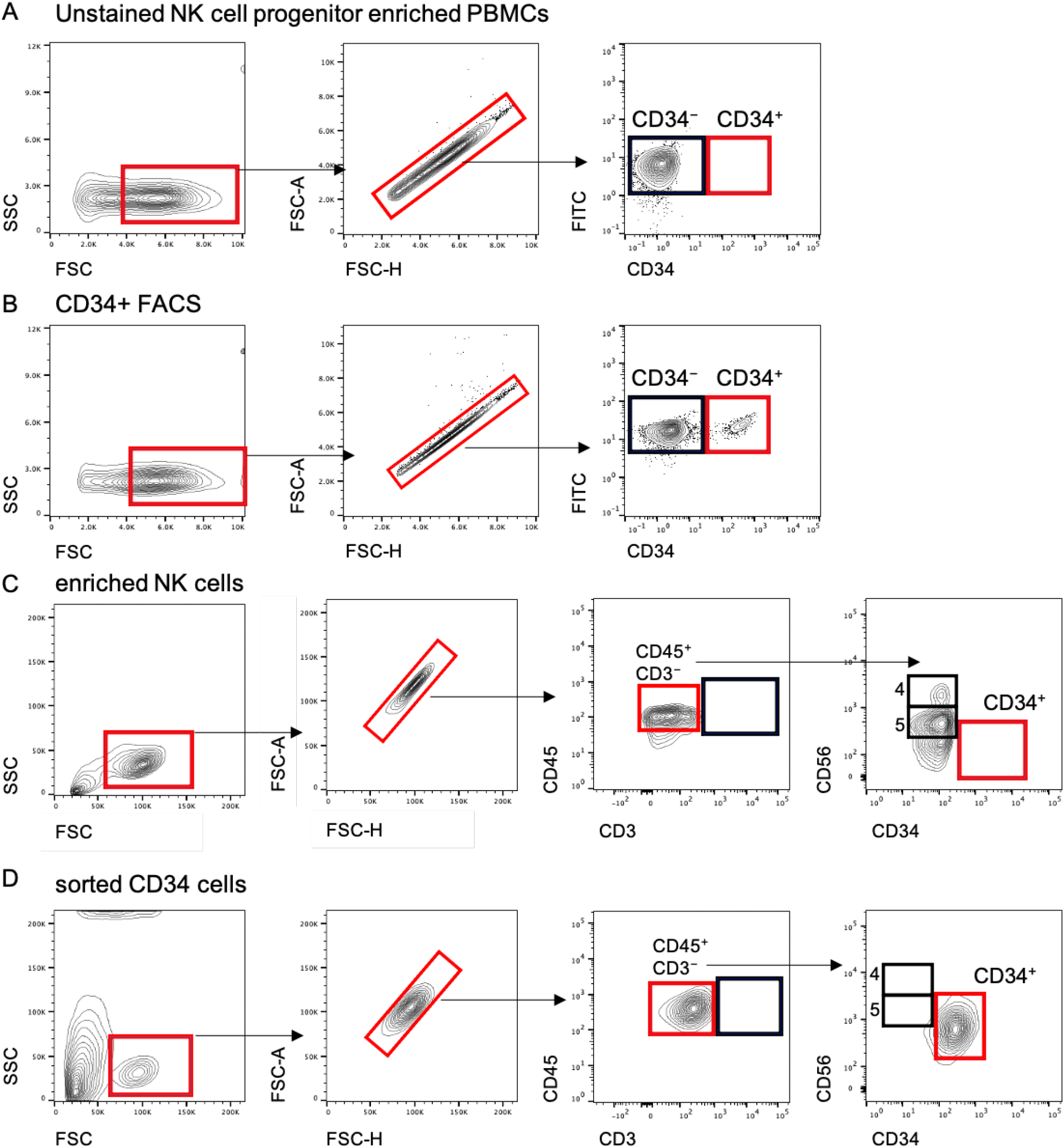
Representative flow cytometric gating strategy for sorting CD34 positive cells from NK cell enriched peripheral blood samples. **A)** Representative unstained enriched NK cell progenitors from peripheral blood mononuclearcells used to set FACS gating for CD34^+^ cell sorting. **B)** FACS sorting of CD34^+^ cells from PE anti-CD34 stained NK enriched peripheral blood mononuclearcells. **C-D)** Representative flow cytometry of enriched NK cells and FACS sorted CD34^+^ cells used for in vitro NK cell differentiation experiments.

**Supplemental Figure 6:**
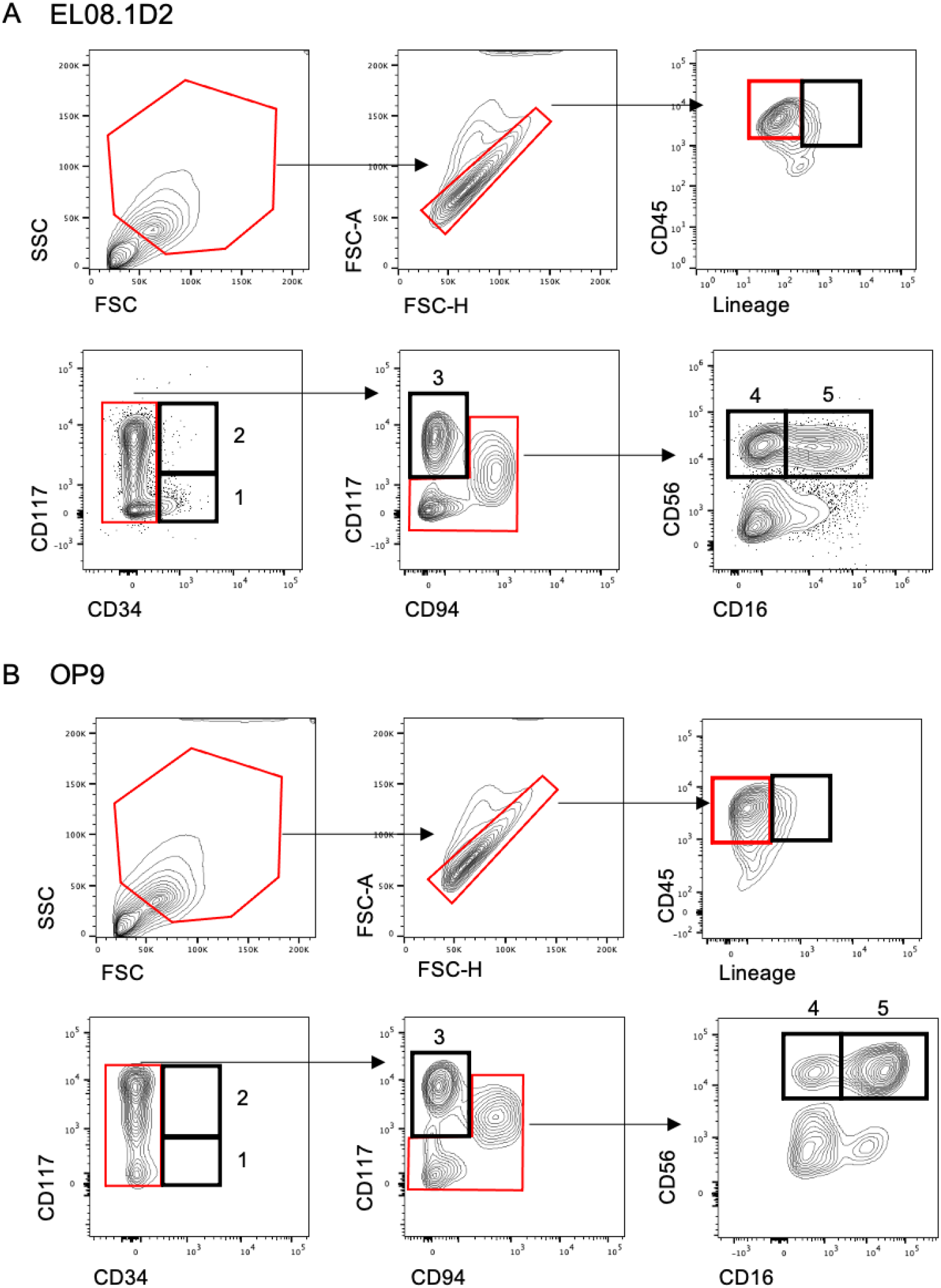
Representative flow cytometric gating strategy for discriminating in vitro NK cell developmental subsets after weeks of culture with either (A) EL08.1D2 or (B) OP9 feeder cells.

**Supplemental Figure 7:**
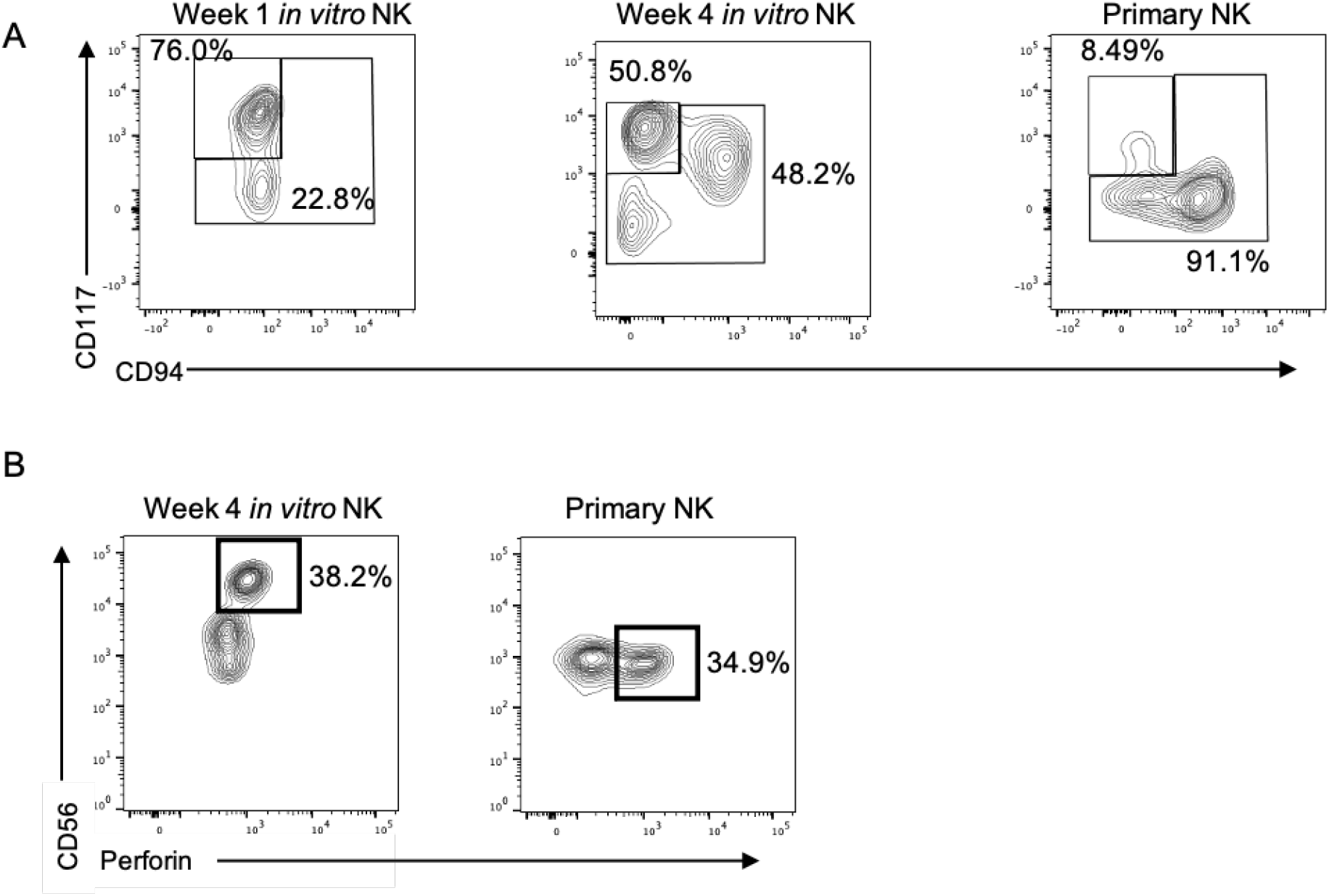
Representative flow plots depicting maturation of in vitro NK cell populations after one and four weeks of culture. After 4 weeks of culture in vitro NK cells become (A) CD94 and (B) perforin positive.

**Supplemental Table 1:**
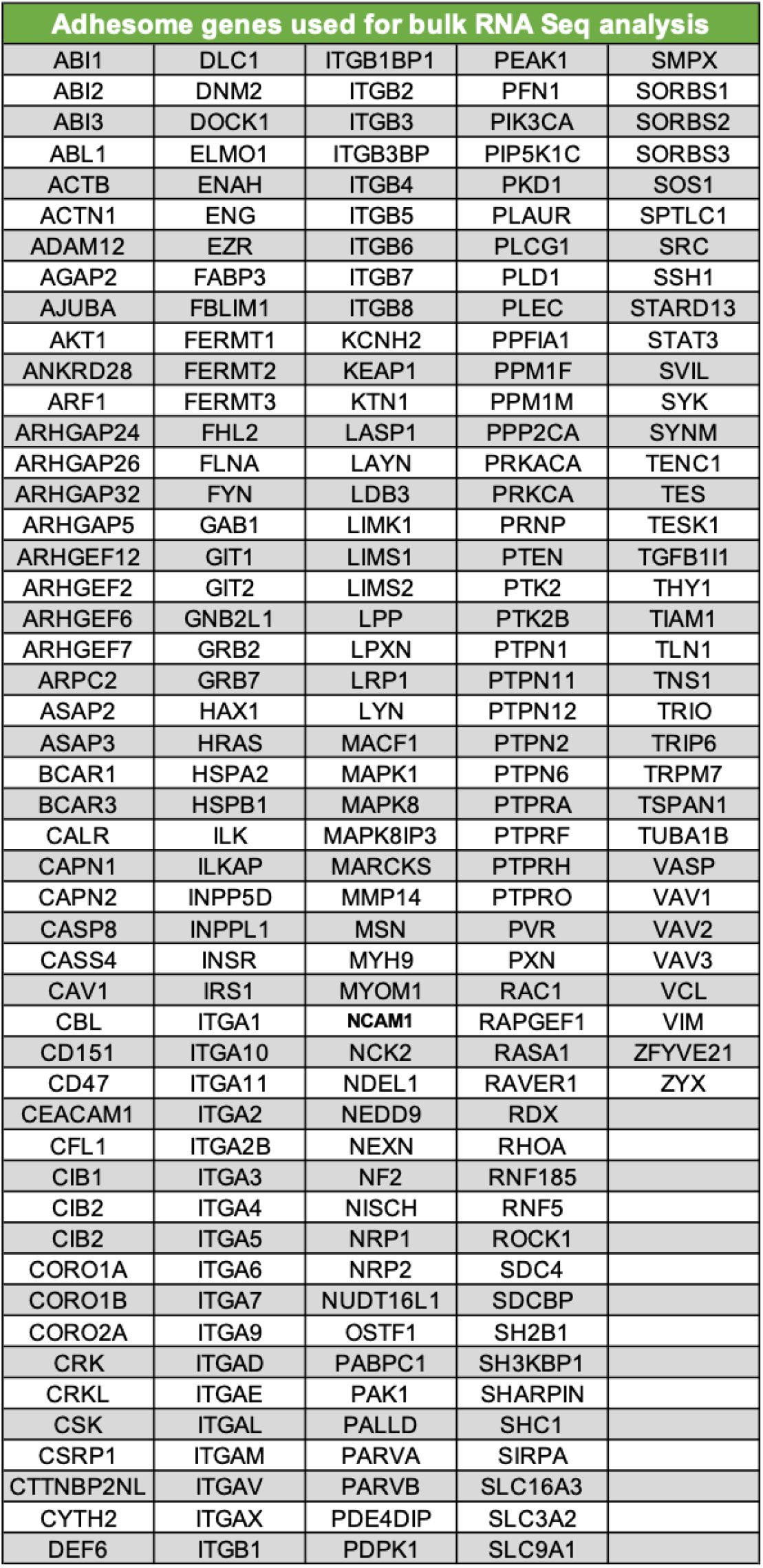
Adhesome gene list for RNA sequencing analysis of peripheral blood and tonsil NK cells. Genes were identified from the consensus integrin adhesome (reference 28).

**Supplemental Table S2: GO Pathway terms (see Excel sheet)**

**Supplemental Table 3:** Adhesome antibody panel for flow cytometric analysis and sorting of NK cells.

| Marker | Fluorophore | Identifier | Source |
| --- | --- | --- | --- |
| CD117 | BV711 | 313230 | Biolegend |
| CD14 | BV421 | 325628 | Biolegend |
| CD16 | PB | 302021 | Biolegend |
| CD16 | PE-CF594 | 562320 | BD Biosciences |
| CD19 | BV421 | 302234 | Biolegend |
| CD3 | BV421 | 344834 | Biolegend |
| CD34 | AF700 | 343526 | Biolegend |
| CD45 | BUV737 | 748719 | BD Biosciences |
| CD56 | BV605 | 318334 | Biolegend |
| CD57 | BV510 | 393313 | Biolegend |
| CD94 | BUV395 | 743954 | BD Biosciences |
| NKp80 | PE-Vio615 | REA845 | Miltenyi Biotec |
| Activated CD29 | FITC | FCMAB389F (HUTS4) | Millipore |
| CD11a/CD18 | APC | 363410 (mAb 24) | Biolegend |
| CD11b | APC-CY7 | 301342 | Biolegend |
| CD11c | BV650 | 301638 | Biolegend |
| CD14 | Viogreen | 130-096-875 | Miltenyi Biotec |
| CD16 | AF700 | 557920 | BD Biosciences |
| CD20 | Viogreen | 130-096-904 | Miltenyi Biotec |
| CD3 | Viogreen | 130-097-582, 130-096-910 | Miltenyi Biotec |
| CD49d | BV785 | 304314 | Biolegend |
| CD49e | APC | 328012 | Biolegend |
| CD56 | BV421 | 562751 | BD Biosciences |
| CD57 | APC-H7 | 555618 | BD Biosciences |
| CD94 | PerCP-Cy5.5 | 562361 | BD Biosciences |
| Human CD45 | BV786 | 563716 | BD Biosciences |
| KIR2D | PE | 130-092-688 | Miltenyi Biotec |
| KIR3DL1/2 | PE | 130-095-205 | Miltenyi Biotec |
| NKp80 | APC | 130-094-845 | Miltenyi Biotec |
| Perforin | PE | 308106 | Biolegend |
| Phalloidin | AF568 | A12380 | ThermoFisher |
| Total CD18 | Pe-Cy7 | 302109 | Biolegend |
| Total CD29 | APC-Cy7 | 303008 | Biolegend |
| Total CD29 | FITC | 303016 (TS2/16) | Biolegend |

**Supplemental Table 4:** Gating strategy utilized for discriminating natural killer cell developmental subsets.

| NKDI Stages | Markers |  |  |  |  |  |  |  |  |
| --- | --- | --- | --- | --- | --- | --- | --- | --- | --- |
|  | CD45 | Lineage | CD34 | CD117 | CD94 | CD56 | NKp80 | CD16 | CD57 |
| Stage 1 | + | - | + | - | - | - | - | - | - |
| Stage 2 | + | - | + | + | - | - | - | - | - |
| Stage 3 | + | - | - | + | - | +/- | - | - | - |
| Stage 4A | + | - | - | +/- | + | ++ | - | - | - |
| Stage 4B | + | - | - | - | + | ++ | + | - | - |
| Stage 5 | + | - | - | - | +/- | + | + | + | - |
| Stage 6 | + | - | - | - | - | + | + | + | + |

**Supplemental Table 5: Complete gene list (see Excel sheet)**

